# Conditional *nmy-1* and *nmy-2* alleles establish that non-muscle myosins are required for late *C. elegans* embryonic elongation

**DOI:** 10.1101/2024.04.12.589286

**Authors:** Kelly Molnar, Shashi Kumar Suman, Jeanne Eichelbrenner, Camille N. Plancke, François B. Robin, Michel Labouesse

**Affiliations:** Laboratoire de Biologie du Développement - UMR7622, Institut de Biologie Paris Seine, Sorbonne Université, 7-9 quai Saint Bernard 75005 Paris, France; Laboratoire Matière et Systèmes Complexes, Université Paris Cité – CNRS UMR7057, 10 Rue Alice Domon et Léonie Duquet - 75013 Paris, France; Development and stem cell program, IGBMC, UMR7104, U964, Université de Strasbourg, 1 rue Laurent Fries, BP10142, 67400 Illkirch, France; CIML - UMR7280, 163 avenue de Luminy, 13288 Marseille Cedex 09, France; Institut Pierre-Gilles de Gennes for Microfluidics, 6 rue Jean Calvin 75005 Paris, France

**Keywords:** *C. elegans*, morphogenesis, non-muscle myosin, mechanotransduction, aggregates

## Abstract

The elongation of *C. elegans* embryos allows examination of mechanical interactions between adjacent tissues. Muscle contractions during late elongation induce the remodelling of epidermal circumferential actin filaments through mechanotransduction. We investigated the possible role of the non-muscle myosins NMY-1 and NMY-2 in this process using *nmy-1* and *nmy-2* thermosensitive alleles. Our findings suggest these myosins act redundantly in late elongation, and that they are involved in the multi-step process of epidermal remodeling. When inactivated, NMY-1 was seen to form protein aggregates.

## INTRODUCTION

Morphogenesis refers to the process by which organisms gradually develop a characteristic 3D form. This can involve an increase in the number of cells, or a change in the configuration of their relative positions, which is accompanied by modifications in their cell-cell membrane adhesions (Lecuit, 2005). Multicellular organisms need to spatiotemporally coordinate the morphogenesis of multiple tissues in time and space (GILMOUR *et al*. 2017; GOODWIN AND NELSON 2021). It has become clear that communication through mechanical inputs plays a key role in ensuring the smooth development of adjacent tissues (AIGOUY *et al*. 2010; ZHANG *et al*. 2011; COLLINET *et al*. 2015; LYE *et al*. 2015). A classical paradigm is that a protein associated with a transmembrane receptor, such as integrin or E-cadherin, undergoes a conformational change that favours the binding of additional proteins. In turn, this can influence protein trafficking, the orientation of planar polarity, junction remodelling, cytoskeleton dynamics, or the translocation of transcription factors to the nucleus, to name a few (DEL RIO *et al*. 2009; AIGOUY *et al*. 2010; LE DUC *et al*. 2010; YONEMURA *et al*. 2010; LEVAYER *et al*. 2011; ZHANG *et al*. 2011; LARDENNOIS *et al*. 2019).

Although, several of the proteins relaying mechanical stress within a cell have been identified, we are far from a detailed understanding of all mechanotransductive pathways (MOORE *et al*. 2010; HU *et al*. 2017; YAP *et al*. 2018; NIETHAMMER 2021). In particular, tissue morphogenesis generally involves repeated mechanical inputs resulting in progressive shape changes (MARTIN *et al*. 2009; SOLON *et al*. 2009; AIGOUY *et al*. 2010; RAUZI *et al*. 2010; ZHANG *et al*. 2011; MAITRE *et al*. 2015; LARDENNOIS *et al*. 2019), but the mechanisms involved in stabilizing cell shapes are only beginning to be discovered. Recent results have emphasized the importance of permanent viscoplastic changes induced by repeated mechanical inputs and the key role of the actomyosin cortex (BONAKDAR *et al*. 2016; DOUBROVINSKI *et al*. 2017; KHALILGHARIBI *et al*. 2019; LARDENNOIS *et al*. 2019; STADDON *et al*. 2019; MOLNAR AND LABOUESSE 2021).

*C. elegans* represents a powerful system to analyze the consequences of mechanical inputs. During *C. elegans* embryonic morphogenesis, an ellipsoidal cell aggregate elongates fourfold into a vermiform shape (Priess & Hirsh, 1986). This occurs in two distinct stages and relies on epidermal cell shape change (Vuong-Brender et al., 2016). The mechanical input comes from the epidermal actomyosin cortex and from the muscles for the early and late stages of elongation, respectively (WILLIAMS AND WATERSTON 1994; PIEKNY *et al*. 2003; GALLY *et al*. 2009; ZHANG *et al*. 2011).

The early stage involves two distinct groups of epidermal cells, the lateral and dorsal/ventral cells. In the lateral cells, there is a greater concentration of non-muscle myosin and a disordered actin network (PIEKNY *et al*. 2003; GALLY *et al*. 2009). Contractions from these two zones flanking the embryo on either side provide the force from the one-fold to roughly 2-fold stages, at which point the muscles become active (VUONG-BRENDER *et al*. 2017). Equally important are the actin filaments in the dorsal/ventral cells, which are arranged circumferentially and make bundles of a few distinct filaments, which provide integrity along the body and cause the force generated by the lateral cells to be directed to the tips of the embryo (VUONG-BRENDER *et al*. 2017).

The late stage begins when the muscles are in place and begin to contract (WILLIAMS AND WATERSTON 1994), at which point the actomyosin cortex in the lateral cells has achieved a circumferential pattern as well (GILLARD *et al*. 2019). Muscles are arranged into four bands just underneath the epidermis to which they are tightly attached, such that muscle contractions induce deformation of epidermal cells (ZHANG AND LABOUESSE 2010; ZHANG *et al*. 2011) . In particular, muscle activity transiently bends the actin bundles beyond a critical angle (Fig. 1A), shown *in vitro* to induce severing by cofilin (MCCULLOUGH *et al*. 2011); in C. *elegans* the severing proteins are villin and gelsolin instead (LARDENNOIS *et al*. 2019). Actin filaments are then restabilized by the p21-activated kinase PAK-1, the alpha-spectrin SPC-1 and the atypical formin FHOD-1 (Fig. 1A) (LARDENNOIS *et al*. 2019). In the absence of PAK-1 and SPC-1, or FHOD-1 and SPC-1, embryos elongate up to the 1.5-fold stage and then regress to their initial lima bean shape (roughly 1.2-fold stage) due to the loss of actin filament integrity (LARDENNOIS *et al*. 2019). Thereby muscles contribute to progressively shorten actin filaments, promote embryo elongation and decrease their diameter.

**Figure 1.**
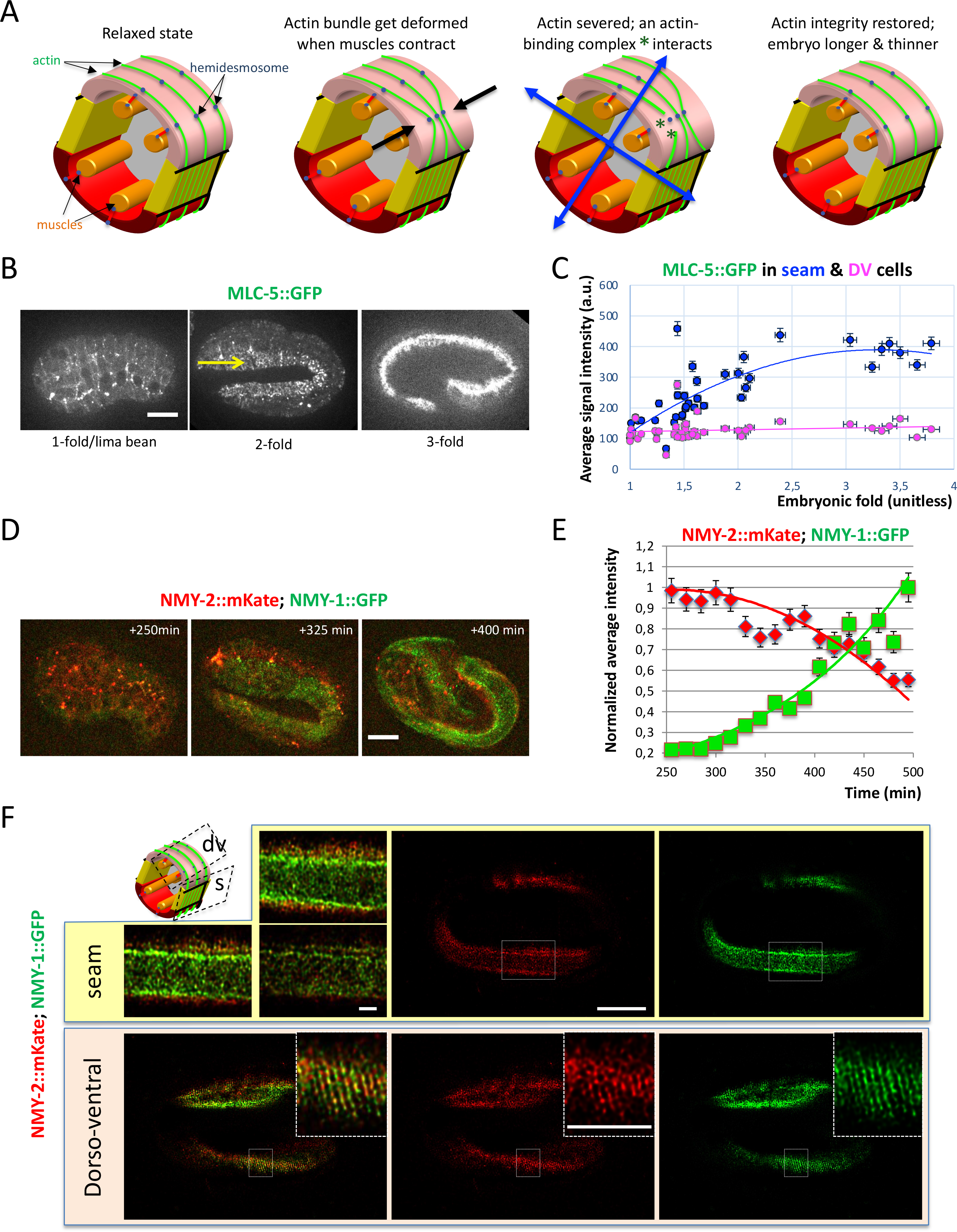
The two non-muscle myosins are present in the epidermis throughout morphogenesis **(A)** Cross-section through an embryo illustrating the main anatomical features of the embryo: the dorsal and ventral epidermis (pink, red), the lateral epidermis (yellow) and the muscles (orange); other tissues not represented for clarity. Muscles are attached to the apical extracellular matrix (not illustrated) surrounding the embryo through hemidesmosomes in the epidermis (blue dots). Actin filaments (green) run circumferentially as bundles. The current model holds that in a typical contraction cycle, muscles will locally displace the actin filaments (black arrows in 2^nd^ drawing), which will trigger their severing (3^rd^ drawing), before they eventually get stabilized by a complex of actin-binding proteins (dark green stars). At the end of the cycle, when the underlying muscles relax, the embryo has become thinner and longer (4^th^ drawing). It is unclear whether the actin-binding complex can hold actin ends to counter the hydrostatic pressure (blue arrows in 3^rd^ drawing) if they drift apart. **(B)** Fluorescence micrographs showing the distribution of the essential myosin light chain MLC-5 (marked by a GFP knockin) at three different stages: lima bean, 2-fold, 3-fold. Note that MLC-5 is enriched in the lateral seam cells (yellow arrow). **(C)** Quantification of MLC-5::GFP fluorescence over time in the lateral and dorso-ventral epidermis. **(D)** Fluorescence micrographs showing the distribution of the two large non-muscle myosin chains NMY-1 (marked by a GFP knockin) and NMY-2 (marked by a mCherry knockin) at different stages; timing starts approximately 125 minutes after the 1-cell stage (black line in E; see video1). **(E)** Quantification of NMY- 1::GFP and NMY-2::mKate fluorescence starting at the lima bean stage. Note the lack of a clear myosin cable along cell contours in the 2-fold embryo (+300 min image). **(F)** Deconvoluted confocal micrographs of NMY-1::GFP and NMY-2::mKate in the lateral seam (yellow box) and dorso-ventral (pink box) epidermis of a 3-fold embryo. Higher magnification of the area marked by a dotted rectangle are shown on the left for seam cells (three consecutive focal planes; 1.75x magnification), or in the top right corner for the dorso-ventral epidermis (4.5x magnification) revealing short circumferential alignments. Scale bars, 10 µm (B, D, F), except 4 µm for the enlargements. Note that NMY-1 and NMY-2 form parallel bands, especially in the dorso-ventral epidermis, but outside the area located above muscles.

The model described above assumes that the ends of severed actin filaments do not get much further apart before they get restabilized by the PAK-1/SPC-1/FHOD-1 complex. However, this might not be the case since hydrostatic pressure exerts a radial force that could be expected to pull those actin ends if they are not held together. Since muscles are arranged orthogonally to actin bundles, they cannot contribute to bring actin ends closer. We thus wondered whether there might be additional players holding actin filaments once severed, and/or bringing them closer before handing them over to PAK-1/SPC-1/FHOD-1 (Fig. 1).

Although spectrins are actin-binding proteins that could fulfil the ascribed function of holding actin ends (LENNE *et al*. 2000; LAW *et al*. 2003; CHOI AND WEIS 2011), we considered the possibility that additional proteins could be required. We focused on non-muscle myosins, because they are obvious motor proteins working on actin. Non-muscle myosins are hetero- hexamers consisting of two myosin heavy chains, two regulatory light chains (MLC-4 in *C. elegans*), which must be phosphorylated to enable motor activity, and two essential light chains (MLC-5 in *C. elegans*) (Vicente-Manzanares et al., 2009). Importantly, several of those hexamers can assemble in a multi-subunit complex of opposite polarity through the C-terminal coiled-coil of the heavy chain. *C. elegans* has two non-muscle heavy chains, NMY-1 and NMY-2. NMY-1 is expressed at the time of elongation, is partially required for elongation and is essential for fertility (PIEKNY *et al*. 2003; KOVACEVIC *et al*. 2013). NMY-2 is expressed in early embryos, and is essential to establish early embryonic polarity and for cytokinesis (GUO AND KEMPHUES 1996; SHELTON *et al*. 1999; MUNRO *et al*. 2004; LIU *et al*. 2010). Both isoforms appear to act redundantly in embryonic morphogenesis, although the precise stage at which they are acting together has not been determined due to the genetic tools used at the time (PIEKNY *et al*. 2003).

To define more precisely how the two heavy chains act, we used temperature sensitive mutants to inactivate NMY-1/-2 at different stages. In addition, we combined them with other players involved in actin dynamics downstream of the muscle-induced actin severing. Our data establish that NMY-1 and NMY-2 are jointly required once muscles become active. We discuss their role compared to spectrins.

## MATERIALS AND METHODS

### Strains and genetic methods

The *C. elegans* control strain N2 and other strains were maintained under standard conditions (Brenner 1974), and were propagated at 20°C unless noted otherwise. A complete list of strains and associated genotypes used in this study are included in Table S1.

### Generation of a *nmy-1* thermosensitive allele

We introduced a mutation in *nmy-1* by CRISPR (Clustered Regularly Interspaced Short Palindromic Repeats) at the position corresponding to the allele *nmy-2(ne3409ts)* affecting changing Leucine-981 to Proline. The corresponding leucine is conserved among non-muscle myosins (see Fig. S1). It was generated using an oligonucleotide repair template (GAAACCGTCCGTGATCTCGAGGAGCAACTCGAGCAAGAtGAACAAGCTAGACAG AAACTGCTTccGGATAAGACGAATGTTGACCAGAGACTTCGAAACCTGGAAGAGCG) carrying two mutations, one to create a non-functional PAM (protospacer adjacent motif) site (AGG to ATG), and one to introduce the desired mutation (TTG to CCG). We co-injected the plasmid encoding Cas9 (CRISPR-associated endonuclease 9) and the sgRNA (single-molecule guide RNA) for *nmy-1* (5’GAGGAGCAACTCGAGCAAG) at 50 ng/µl, the *nmy-1* oligonucleotide repair template at 20 ng/µl, along with the plasmids pRF4 and pBSK2 each at 100 ng/µl in the strain ML2540 (10.7554/eLife.23866) carrying a CRISPR-generated NMY- 1::GFP fusion, so as to track the putative mutant protein. We subsequently picked 88 Roller animals and screened by PCR for the presence of the mutation using the primers 5’TCAAGCTCACCGCTTTAATTATGAAC and 5’CCCATTTTCTCGGCCAAGTGATCT to find one positive hit that could be recovered, which we named *nmy-1(mc90ts)*. The allele *nmy-1(mc90ts)* had two mutations, as verified by sequencing the *nmy-1* locus from homozygous animals, one corresponding to the desired L>P change and one corresponding to the PAM mutation, such that the protein sequence 959-QEEQARQKLLL became 959- QDEQARQKLLP. Homozygous *nmy-1(mc90ts)* animals were sterile at 25°C but not at 15°C (see results). We assume that thermosensitivity was due to the Leu to Pro change rather than to the upstream synonymous Glu to Asp change at the PAM site in the repair template.

### RNA interference (RNAi)

RNAi experiments were performed by feeding on *HT115 Escherichia coli* bacteria strains generating double-stranded RNA (dsRNA) from the Ahringer-MRC feeding RNA interference (RNAi) library (Kamath et al., 2003). RNAi feeding was performed using standard procedures, with 100 μg ml^−1^ ampicillin/1 mM IPTG (Sigma). Empty L4440 RNAi vector served as a control.

### Hatch count and arrest count protocols

Strains carrying a theremosensitive mutation were stored at 15°C on agar plates. Embryos at different developmental stages on the plates were removed with an eyelash tool, washed in M9, and put on a 5% agarose pad with M9 on a slide sealed in paraffin oil. The slide was then imaged using a Roper Scientific spinning disk system with an immersion oil 40x objective (Zeiss Axio Observer Z1 microscope, Yokogawa CSUX1-A1 spinning disk confocal head, Photometrics Evolve 512 EMCCD camera, Metamorph software). Images were initially taken at 20°C until the incubation chamber attached to the microscope was set to 25°C, and imaging was continued overnight (8 hrs). A count was then performed to determine how many embryos of a particular initial developmental stage could make it to hatching, and at what stage they had arrested, if so.

### Fluorescence microscopy

Spinning disk fluorescence imaging was performed with a 63× or 100× oil-immersion objective, NA=1.4. The temperature of the microscopy room was maintained at 20°C. Images of embryos were acquired using time-lapse mode with a 110 ms exposure at intervals depending on the experiment. Laser power and exposure times were kept constant throughout the experiments for specific strains and their controls. Images of Fig. 1F were acquired with a LSM 980 Airyscan2 Zeiss confocal system equipped with 488 nm and 561 nm excitation laser lines and an oil immersion objective with NA=1.4. We performed a simultaneous acquisition for the green and red channels in order to get both stainings in the same time in confocal mode. The images were deconvolved with the Huygens software.

### Image processing and quantification

All images were treated by first subtracting the background with a rolling ball radius of 50 pixel. Any stacks were projected using maximum intensity. In Fig. 4, the average displacement was obtained by tracking stable points of high intensity in the epidermis and calculating the displacement using the following equation:

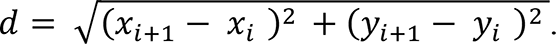

The images in Fig. 4 were three-plane projections from stacks taken with an inter-plane distance of 0.8 μm to capture the epidermis. The distribution of the signal was obtained by measuring the mean intensity and standard deviation of a square of 45X33 pixels inside the H1 cell of the lateral epidermis, distribution = σ/. Images with an even distribution of signal (meaning no bright spots) have a low standard deviation σ, and therefore a low distribution. In the case of bright spots against a black background, the value of σ increases, and therefore so will the distribution.

## RESULTS and DISCUSSION

### The concentration of non-muscle myosins increases until the 2-fold stage

To better understand when non-muscle myosins are required during elongation, we examined the fluorescence of three CRISPR knockin strains marking the essential light chain MLC-5, NMY-2 and NMY-1. We imaged randomly picked MLC-5::GFP embryos and examined both their stage and the GFP fluorescence level. We observed that the fluorescence increased between the one- and 2-fold stages, and was followed by a slight plateau in the lateral cells, and a mostly steady value in the DV cells (Fig. 1B-C). Using an NMY-2::mKate; NMY-1::GFP double knockin strain, we observed that only NMY-2 started to decline after the beginning of lima bean stage, whereas NMY-1 increased after that stage (Fig. 1D-E). At higher magnification, we observed that NMY-2 was faint with no clear pattern in the lateral cells but formed aligned puncta in DV cells, whereas NMY-1 was enriched in the lateral epidermis cells as aligned puncta and colocalized in DV epidermal cells with NMY-2 puncta (Fig. 1F). These results indicate that the myosin population is not static. Furthermore, they are compatible with the possibility that the two non-muscle myosin motors could be acting redundantly with one another during late elongation.

### Inhibition of NMY-1 and NMY-2 arrests late elongation

To test the functions of the two non-muscle myosin heavy chains, we used conditional alleles. A temperature sensitive *nmy-2* allele had previously been described, *nmy-2(ne3409ts)*, which changes a conserved Leucine residue among heavy chains into a Proline, NMY-2(L981P) (LIU *et al*. 2010). We engineered an NMY-1(L970P) change by CRISPR at the position homologous to that of *nmy-2(ne3409)*; the strategy also involved a change of E960 into D (see Methods and Fig. S1). The resulting allele, named *nmy-1(mc90ts)*, induced a thermosensitive sterile phenotype but very little lethality (Table 1; Fig. 2A,2B-line 4). We assume that the sterility was due to the L970P mutation rather than to the synonymous E960D change, because the *nmy- 2(ne3409ts)* at the homologous position is conditional, but cannot exclude additive effects of both changes. At the non-permissive temperature, the allele *nmy-1(mc90ts)* behaved like the previously known *nmy-1* missense allele *sb113*, which induced <10% lethality and partial sterility, but was not as severe as the presumptive null allele *sb115*, which induced over 50% embryonic and larval lethality (PIEKNY *et al*. 2003). Since the mutant protein NMY-2(ne3409ts) unfolds and becomes very rapidly inactive when animals are shifted to 25°C (LIU *et al*. 2010), we expected the mutant protein NMY-1(mc90ts) to behave likewise upon a temperature shift.

**Table 1**
| Genotype | 20°C | 25°C |
| --- | --- | --- |
| wild-type | 556 | 651 |
| *nmy-1(mc90ts)* | 26 | 5 |
| *nmy-2(ne3409ts)* | 463 | 261 |
| *nmy-2(ne3409); nmy-1(mc90ts)* | 8 | 0 |
| wild-type | 55,6 | 65,1 |
| *nmy-1(mc90ts)* | 2,9 | 0,5 |
| *nmy-2(ne3409ts)* | 46,3 | 26,1 |
| *nmy-2(ne3409); nmy-1(mc90ts)* | 0,8 | 0 |

**Figure 2.**
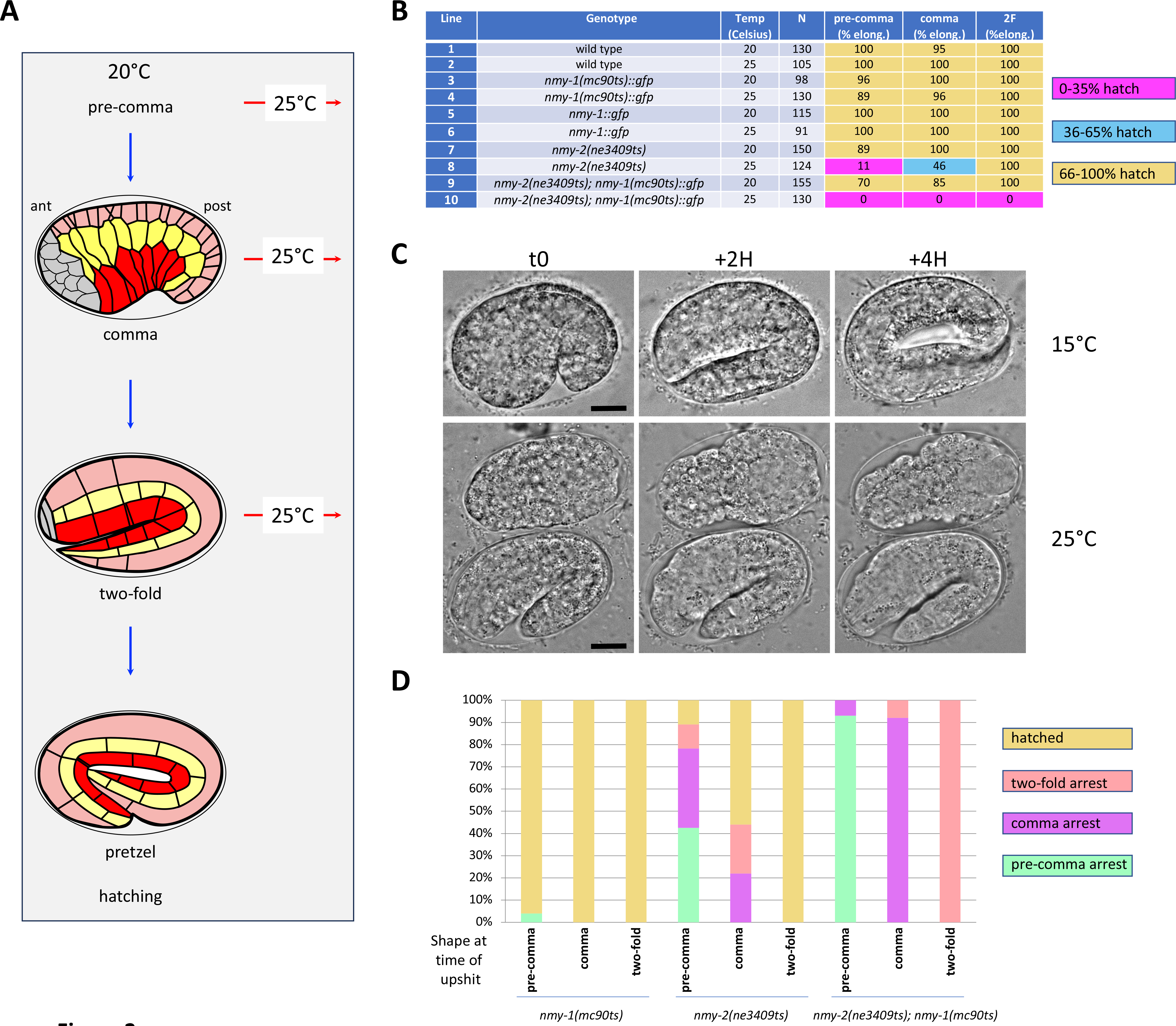
NMY-1 and NMY-2 are required during the morphogenetic phase driven by muscles **(A)** Outline of the temperature shift experiments: embryos were maintained at 20°C and shifted to 25°C at different stages (prior to comma, comma, or two-fold). **(B)** Percentage of hatching when embryos were left at 20°C or shifted to 25°C at the stage indicated in the top row. **(C)** DIC images of *nmy-2(ne3409ts); nmy-1(mc90ts)* left at 15°C (top row) or shifted to 25°C (bottom row). Note that both lima bean and 1.8-fold embryos essentially did not elongate any further after the shift to 25°C at t0; scale bar 10 µm. **(D)** Percentage and arrest stage of single *nmy-2(ne3409ts)*, *nmy-1(mc90ts)* and double mutants shifted to 25°C at the stage indicated on the X axis. Note that for the double mutant, all 2-fold embryos arrested at the 2-fold stage.

Using these two conditional alleles, we investigated at which stage they are required during elongation by raising mutant embryos at 20°C and then shifting them to the non-permissive temperature (Fig. 2A). Homozygous *nmy-2(ne3409ts)* embryos displayed embryonic lethality when shifted at an early elongation stage, however they could elongate if shifted at or beyond the 2-fold stage (Fig. 2B**-**line8). By contrast, double *nmy-2(ne3409ts); nmy-1(mc90ts)* mutants displayed 100% embryonic lethality irrespective of the time at which they were shifted to 25°C (Fig. 2B-line10), but could in general hatch if maintained at 20°C (Fig. 2B-line9).

We further examined at which elongation stage non-muscle myosin mutants arrested. When *nmy-2(ne3409ts)* embryos were shifted to 25°C at the pre-comma stage, about 40% remained pre-comma and 40% made it to the comma stage (Fig. 2D). When shifted at the comma stage, 50% *nmy-2(ne3409ts)* progressed to the 2-fold stage. When double *nmy-2(ne3409ts); nmy- 1(mc90ts)* mutants were shifted to 25°C, they immediately stopped elongation at the stage at which the temperature had been raised (Fig. 2C-D). Importantly, they did not regress to an earlier body morphology as we had observed in *spc-1 pak-1* or *fhod-1; spc-1* double mutants (Fig. 2C). These results show that NMY-2 is required during elongation, but not when muscles become required. Moreover, we conclude that NMY-1 and NMY-2 are continuously required for elongation at all stages, that they act redundantly, but that they are not required to maintain body shape. These observations are also consistent with the notion that both non-myosin mutant proteins become very rapidly inactive at 25°C. The arrest phenotype observed upon an early elongation shift resembles that observed in strong *let-502, mlc-4* or *mlc-5* deficient embryos, as had previously been reported (PIEKNY *et al*. 2003). However, in contrast to the arrest phenotype observed when *nmy-2(ne3409ts); nmy-1(mc90ts)* are shifted to 25°C at the 2-fold stage, *let- 502(sb92ts)* embryos shifted to 25°C at the 2-fold stage did not arrest during elongation (DIOGON *et al*. 2007). One possibility could be that another kinase such as PAK-1 or MRCK-1 acts in parallel to LET-502/Rho-kinase to phosphorylate MLC-4/RMLC at that stage (GALLY *et al*. 2009).

Our results confirm and much extend an earlier study suggesting that the two non-muscle myosins NMY-1 and NMY-2 are redundantly required during embryonic elongation (PIEKNY *et al*. 2003). Those previous experiments involved RNA interference against *nmy-2* in the background of the non-conditional and putative null alleles *nmy-1(sb113)* or *nmy-1(sb115)*. As RNAi against *nmy-2* had to be mild to allow cytokinesis, it did not enable the authors to define when NMY-1 and NMY-2 act during elongation. Our results establish that neither NMY-1 nor NMY-2 alone is required, but that they are constantly and redundantly required during late embryonic elongation, which depends on muscle input. Moreover, we conclude that NMY-2 has a more critical role during early elongation since all single *nmy-2* mutants, but only a few single *nmy-1* mutants, arrested when shifted to 25°C during early elongation. Consistent with these results, NMY-1, NMY-2, and the myosin essential light chain MLC-5 are expressed in the epidermis throughout elongation.

### Elongation of *nmy-2 (ne3409ts); nmy-1(mc90ts)* resumes after arrest

As reported above, the double *nmy-2 (ne3409ts); nmy-1(mc90ts)* mutant always arrested during elongation, regardless of the initial stage of the embryos at the time of the temperature upshift. Nevertheless, these embryos could maintain normal muscle activity for several hours after the temperature shift. We thus wondered whether elongation could resume after an arrest. To test this possibility, we shifted *nmy-2 (ne3409ts); nmy-1(mc90ts)* embryos at 25°C for 45 min, using the incubator on the microscope to control the temperature, then then back to 15°C for 5hrs in a different incubator before imaging. As described in Fig. 3A, 60% of 2-fold *nmy-2 (ne3409ts); nmy-1(mc90ts)* embryos were able to hatch, as compared to 0% when left at 25°C. The resumption of elongation after arrest suggests that the temporary absence of NMY-1 and NMY-2 did not permanently damage any of the molecular-level components involved in this developmental stage. Finally, we investigated for how long could elongation be paused and then successfully restarted. To test this, we progressively increased the duration of the pause and found that the curve was sigmoidal, with a mid-height value of about 2,5 hours, and that it levelled off around 5 hours (Fig. 3B).

**Figure 3.**
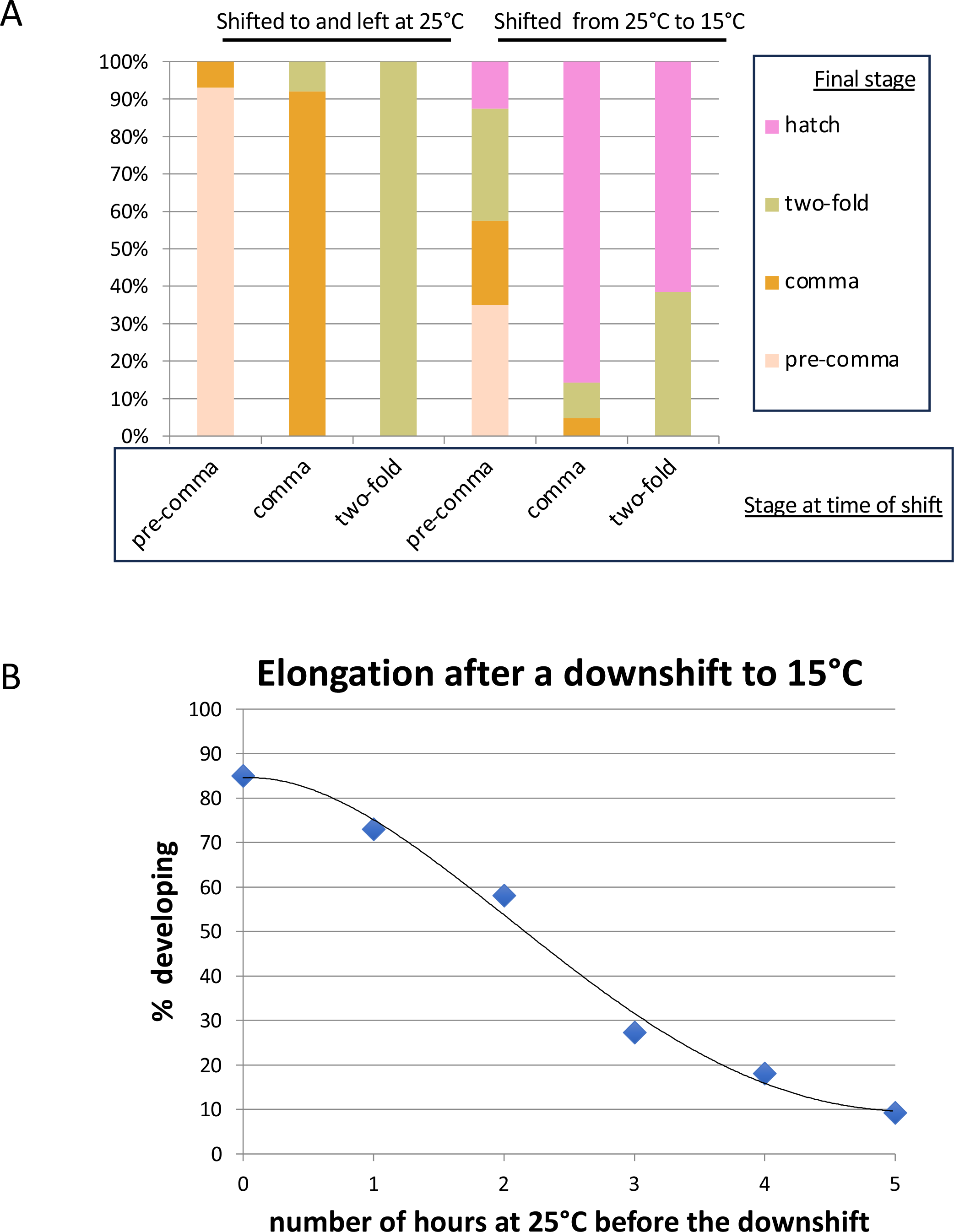
Resumption of elongation. **(A)** Double *nmy-2(ne3409ts); nmy-1(mc90ts)* mutants were shifted to 25°C at the stage indicated at the bottom and left overnight at 25°C (left three columns, taken from Fig. 2D) or left at 25°C for 1 hour, causing arrest, then shifted for 5 hours at 15°C (right three columns). Note that elongation could resume in many cases. **(B)** Quantification of the percentage of *nmy- 2(ne3409ts); nmy-1(mc90ts)* embryos that could resume elongation after being left at 25°C for various duration and then shifted back to 15°C (N>500). The polynomial fit probability is 0.9903.

### *nmy-2 (ne3409ts); nmy-1(mc90ts)* has normal muscle activity at 25°C

We considered two explanations to account for the fact that *nmy-2(ne3409ts); nmy-1(mc90ts)* embryos stop elongation when shifted to 25°C at the 2-fold stage. One possibility would be that these non-muscle myosins are required in muscles, for instance that in their absence muscles do not contract strongly enough or do not transmit properly the contractions to initiate actin filament bending and fail to trigger their reorganisation. Another possibility would be that they act in the epidermis to help reorganise actin filaments.

To test these options, we examined whether *nmy-2(ne3409ts); nmy-1(mc90ts)* embryos twitch normally, taking advantage of the fact that the *nmy-1(mc90ts)* allele is marked by GFP.

Specifically, we used irregularities in the NMY-1::GFP pattern to monitor the twitching pattern of control *nmy-1::gfp* embryos, of embryos in which muscles had been made inactive by RNAi treatment against *unc-112* which is essential to assemble myofilaments (ROGALSKI *et al*. 2000), and of *nmy-2(ne3409ts); nmy-1(mc90ts)* double mutants.

Analysis of NMY-1::GFP videograms showed that embryonic movements can be decomposed into two fundamental movements: rotation and twitching. The first was to follow the rotations of the cylindrical body within the eggshell of 90°, which is triggered by muscle activity (YANG 2017). The second was measured by tracking landmarks in the lateral epidermal cells in-between body rotations (Fig. 4A, lower panels). We found that both control and double non-muscle myosin mutants at 25°C could rotate on average every 12 seconds (Fig. 4A-B upper panels; Fig. 4D), and contract locally over 0.73 ± 0.06 µm/sec, as compared to the control *nmy- 1::gfp*, 0.81 ± 0.06 µm/sec when placed at 25°C (Fig. 4E). By contrast the muscle deficient strain exhibited no rotations and contracted 0.21 ± 0.01 µm/sec (Fig. 4C-E). We conclude that the double mutant is not lacking the mechanical input from the muscles needed to drive elongation. Furthermore, because we measured the behaviour of the epidermis, the muscles are also properly transmitting forces to the neighbouring tissue.

The results described above thus suggest that both non-muscle myosins are required in the epidermis. Which function could non-muscle myosins perform in the epidermis? As recalled above (see introduction), we previously suggested based on genetic and imaging data that the repeated muscle contractions induce the severing and shortening of circumferential actin filaments (LARDENNOIS *et al*. 2019). We could not at the time define what happens with severed actin ends. One possibility is that the hydrostatic pressure building up during elongation could be expected to pull them apart. Although spectrin may keep actin filaments together, since spectrins are considered as springs and ß-spectrin can bind actin (LENNE *et al*. 2000; LAW *et al*. 2003; CHOI AND WEIS 2011), our data posit non-muscle myosins as perfect candidates to keep actin ends together.

Non-muscle myosins have two well-characterized activities, actin binding and actin pulling through their power stroke, which requires myosin regulatory light chain (MLC-4 in *C. elegans*) phosphorylation. We considered the possibility that muscle contractions could locally activate MLC-4 and attempted to test this idea by examining a wild-type MLC-4::GFP marker in control embryos, but failed to record any obvious such event (data not shown). We did not directly test in which cells NMY-1 and NMY-2 are required. However, we note that when wild-type MLC- 4 is expressed under a dorso-ventral epidermal promoter in *mlc-4* null mutant embryos, the embryos elongate up to the 2.5-fold but do not reach full elongation (GALLY *et al*. 2009), indicating that MLC-4 activity, and hence NMY-1/2 activity, is most likely also required in dorso-ventral cells. In part because the precise organization and polarity of actin filaments within dorso-ventral actin bundles is unknown (COSTA *et al*. 1997), we cannot currently determine whether non-muscle myosins are only required to maintain actin ends together or if they pull on severed actin filaments to bring them closer before FHOD-1 activity. If actin filaments all have the same polarity, it would tend to exclude a pulling function, since pulling of actin filaments by non-muscle myosin relies on filaments of opposite polarity. One strategy to discriminate between both models would be to introduce a secondary mutation in NMY-1 preventing ATP binding (OSORIO *et al*. 2019), which is beyond the scope of this study.

### Inactivated NMY-1 forms aggregates

During the course of the experiments described above, we noticed that the NMY- 1(mc90ts)::GFP protein aggregated in the epidermis at the non-permissive temperature (Fig. 5A). The distribution of these aggregates was quantified in the lateral cells (see Methods), whereby a large value of the ratio between the standard deviation σ over the mean intensity () indicated a less uniform distribution. Using this approach, we measured a value of 0.5 ± 0.1 at 25°C for control NMY-1::GFP embryos, the most uniform distribution of all the strains.

**Figure 4.**
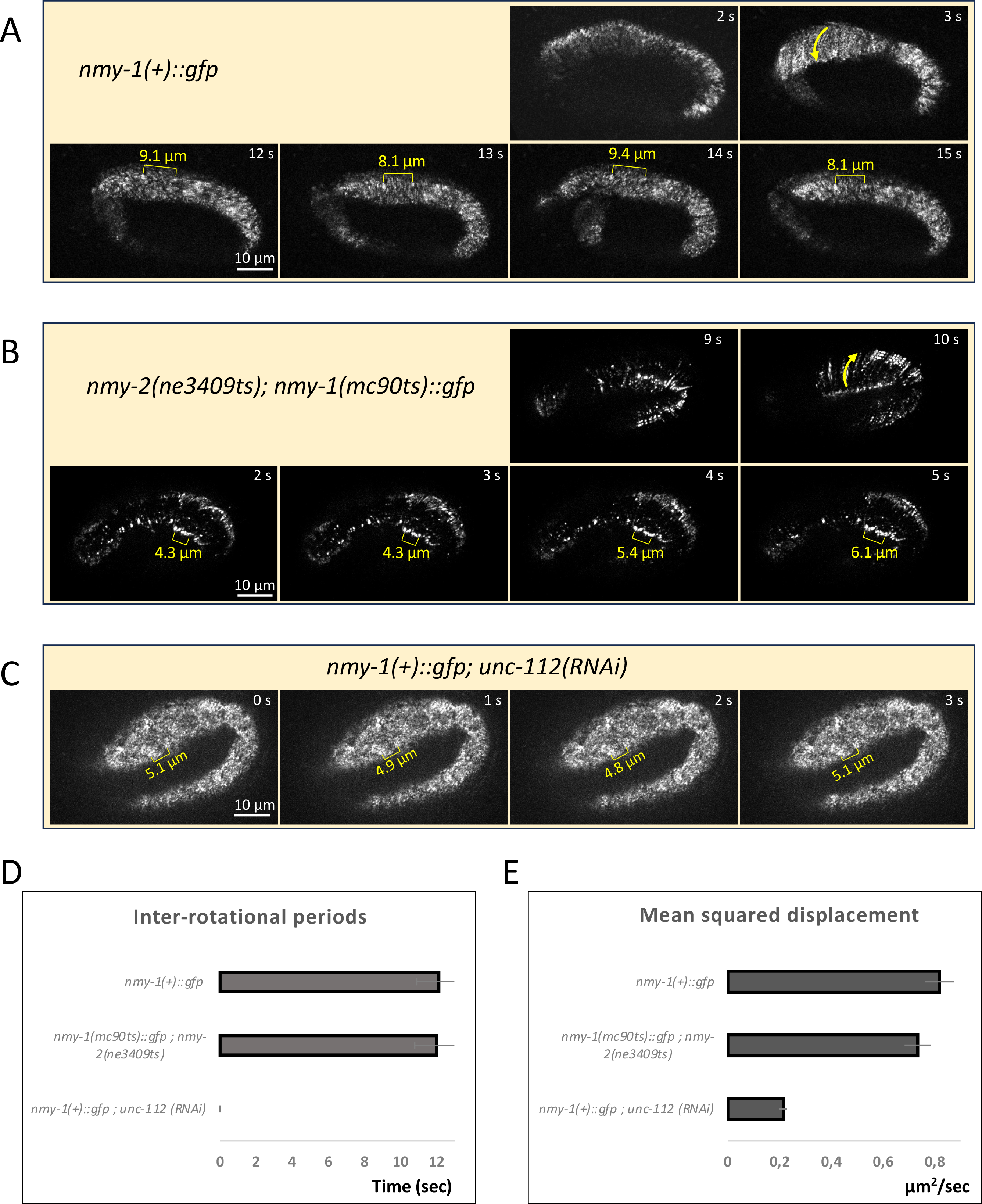
Depletion of NMY-1 and NMY-2 does not affect muscle contractions **(A)** Spinning disk micrographs from Video2 illustrating a rotation (top two images; see yellow arrow) or local contractions (bottom four images) in *nmy-1[mc82(nmy-1::gfp)]* knockin embryos grown at 25°C; timing refers to the video. Note in the bottom four images the local magnitude of contraction between two brighter NMY-1::GFP points (yellow brackets with their size above). **(B)** Spinning disk micrographs from Video3 illustrating a rotation (top two images; see yellow arrow) or local contractions in double *nmy-2(ne3409ts); nmy-1(mc90ts)::gfp* shifted to 25°C when they reached the 2-fold stage; timing refers to the video. Note the local contractions (yellow brackets with their size above). **(C)** Spinning disk micrographs from Video4 the local contractions in a *nmy-1[mc82(nmy-1::gfp)]* embryo treated by RNAi against the gene *unc-112* and raised at 25°C; timing refers to the video. Note the local contractions (yellow brackets with their size above). **(D)** Quantification of the time elapsed between embryo rotations, defined as at least 90° about the centre line of the embryo, as observed in videos similar to those shown in (A-C); *unc-112(RNAi)* failed to rotate. **(E)** Quantification of bright NMY-1::GFP particles lateral movements consecutive to muscle twitching measured from videos such as those in (A-C). The number of embryos was at least 18 for each genotype in all quantifications.

**Figure 5.**
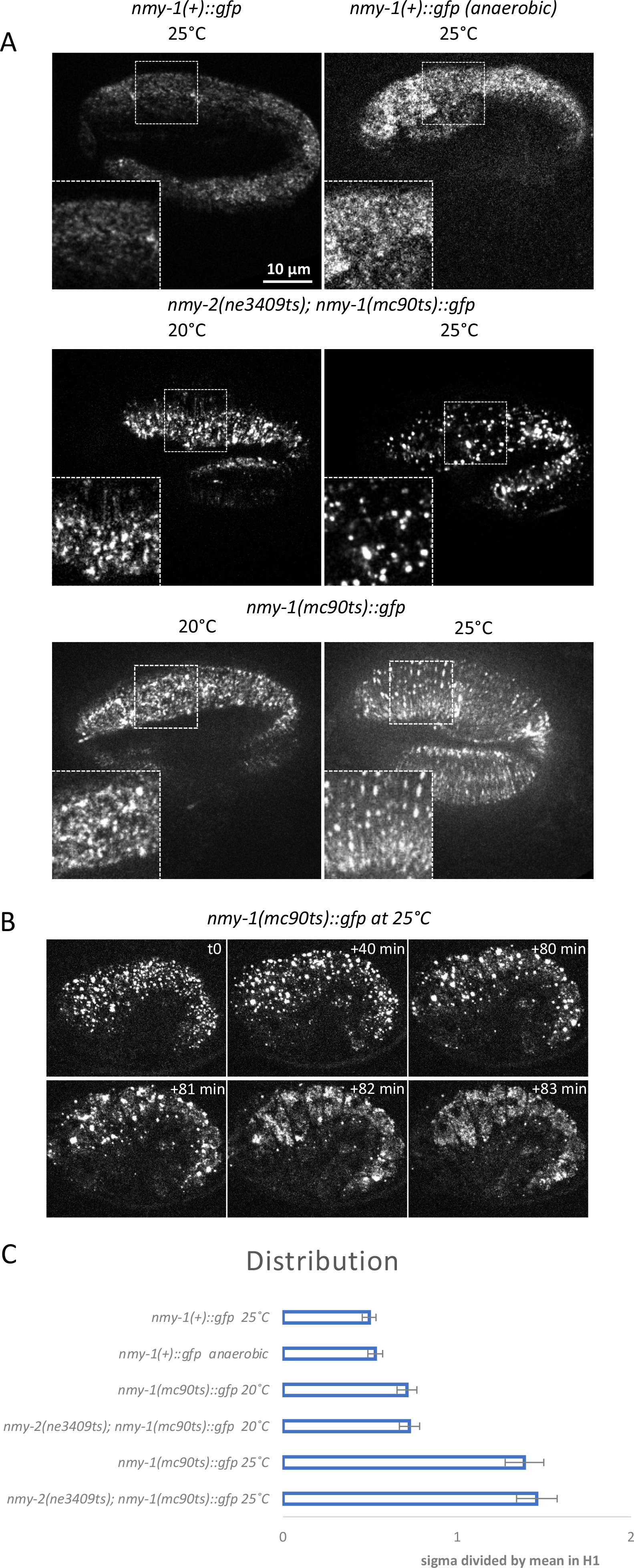
NMY-1(mc90ts)::GFP spots aggregate at the non-permissive temperature **(A)** Spinning disk micrographs illustrating NMY-1::GFP particles in different genotypes when embryos were raised at 20°C, shifted to 25°C after reaching the 2-fold stage, or grown at 25°C in anaerobic conditions. Enlargements (1.75x) of the boxed areas are shown in the bottom right corner. **(B)** Spinning disk micrographs illustrating the further aggregation of NMY-1(mc90ts)::GFP particles, until their rapid dissolution; images taken from Video6. **(C)** Quantification of the distribution of NMY-1::GFP particles brightness measured in the H1 seam cell and expressed as standard deviation of the signal divided by the mean. The number of embryos was at least 25 for each genotype.

It is noteworthy, that the NMY-1(mc90ts) displayed some degree of aggregation even at 20°C as compared to control NMY-1::GFP (see Fig. 5A). At 25°C, we observed a marked aggregation for the single mutant *nmy-1(mc90ts)* (1.4 ± 0.1) as well as for the double mutant *nmy-2(ne3409ts); nmy-1(mc90ts)* (1.5 ± 0.1) (Fig. 5A-B). Interestingly, oxygen depletion, which has a marked effect on the regulatory light chain MLC-4 distribution (GALLY *et al*. 2009), did not modify that of wild-type NMY-1::GFP. It implies that the aggregation of NMY- 1(mc90ts) does not result from ATP-depletion, consistent with the fact that the *mc90* mutation is located in the coiled coil region of NMY-1 (Fig. S1).

We also investigated the long-term fate of these aggregates and found that after the initial inactivation, the many small clusters began to fuse for roughly 1.5 hours, until a sudden transition occurred within 3 minutes. At that point, the aggregates disappeared in a highly coordinated fashion. Note that the images used for the quantification of the distribution of the aggregates were taken on average one hour after the temperature shift had occurred, and thus correspond to roughly the half-point in the time-evolution of the particles. Our findings are consistent with the literature, indicating that non-functional non-muscle myosins in both *C. elegans* oocytes, and human thrombocytes tend to aggregate (ALTHAUS AND GREINACHER 2009; SUN *et al*. 2020).

## Conclusion

Altogether, our data identify the presence of both NMY-1 and NMY-2 in the epidermis during late elongation. We found that either NMY-1 or NMY-2 can support late elongation but that if both are absent elongation beyond the 2-fold stage failed. Not only did their combined absence cause a hatching failure, but the arrest in development was immediate, which is also the case for earlier developmental stages in the embryo. Importantly, this arrest could not be attributed to less muscle movement, and thus to abnormal mechanotransduction in the epidermis. We found that the arrest could be reversed by returning the embryos to < 20°C, indicating that the absence of both non-muscle myosin heavy chains did not permanently affect actin integrity, nor any other epidermal structure. Our data are compatible with the possibility that this myosin pair acts to reduce the length of the epidermal actin filaments by holding or pulling the severed ends together.

## Data Availability Statement

Strains and plasmids are available upon request. The authors affirm that all data necessary for confirming the conclusions of the article are present within the article, figures, and tables.

## Acknowledgements

The authors greatly acknowledge France Lam from the IBPS Imaging Facility for running the Zeiss Airyscan confocal microscope and for performing deconvolution of the images. The IBPS Imaging facility is supported by Region-Île-de-France, Sorbonne-University and CNRS. This work was supported by an Agence Nationale de la Recherche grant to Michel Labouesse (project number ANR-18-CE13-0008-01).

## Supplementary figures

**Supplementary figure 1.**
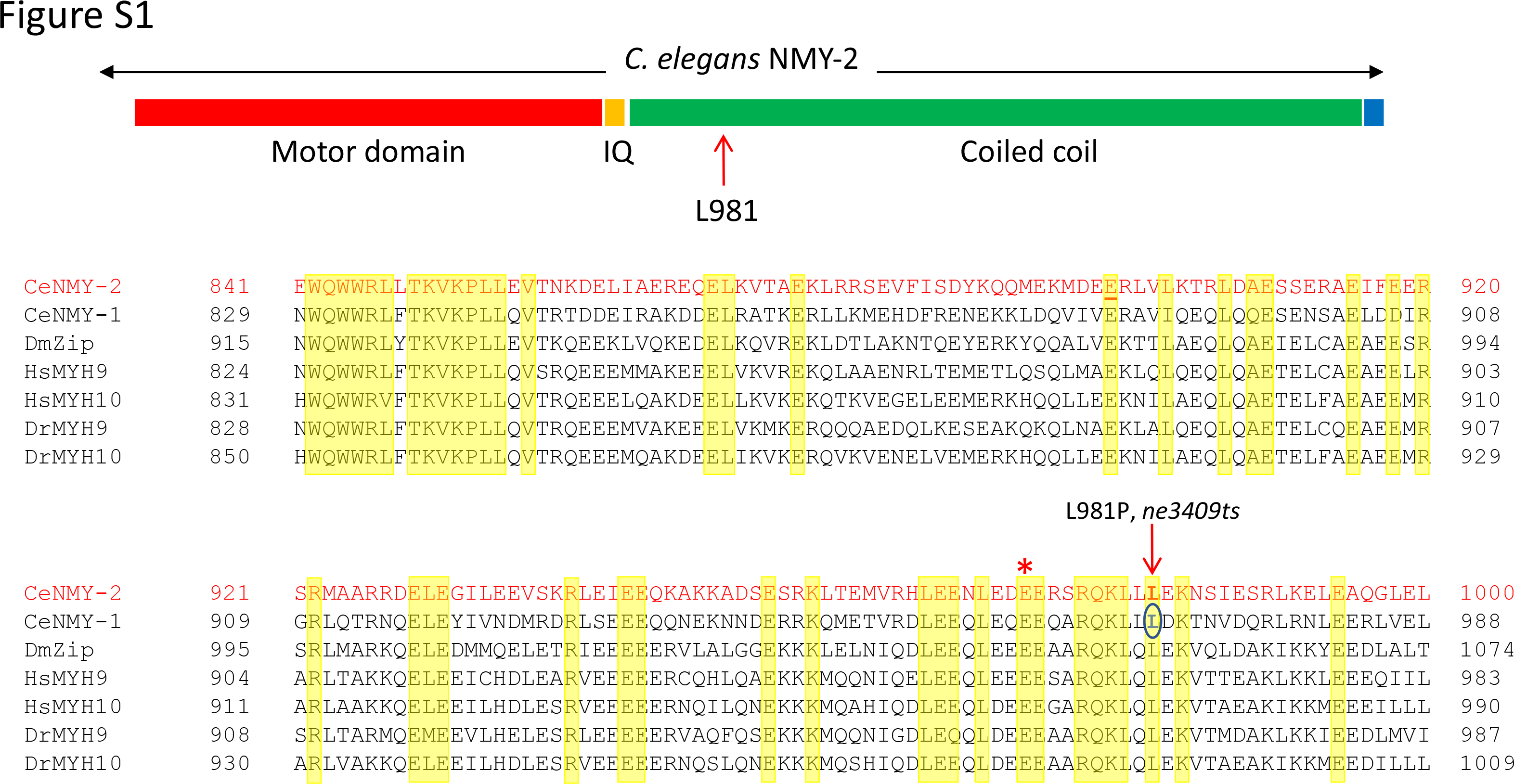
Sequence alignment of non-muscle heavy chains and positions of the *nmy-1* and *nmy-2* conditional alleles. Upper bar, schematized structure of a non-muscle heavy chain, indicating the position of the residue L981 which is mutated to a proline in *nmy-2(ne3409ts)*. Below, alignment of seven heavy chains starting in the IQ region. The residue altered in *nmy-2(ne3409ts)* (red arrow) is conserved and was also mutated in *nmy-1(mc90ts)* (green circle). Red asterisk, position of the secondary mutation introduced to mutate the PAM site (959-QEEQARQKLLL to 959- Q**D**EQARQKLLL).

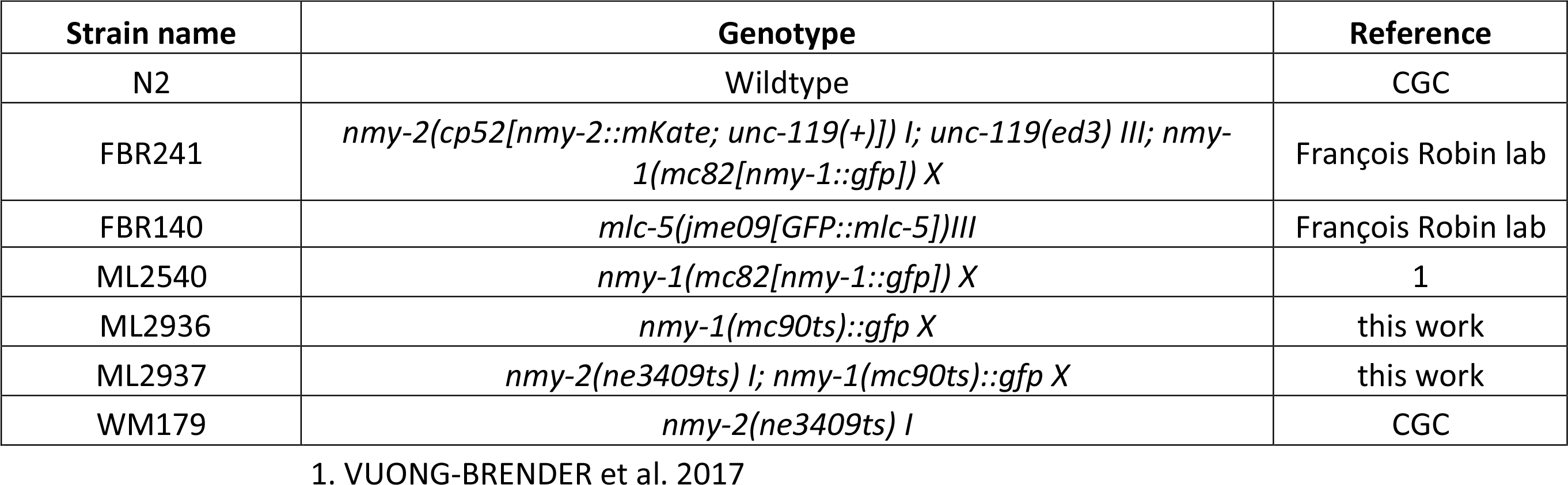

## Supplementary videos

**Video1**: Temporal distributions of a strain carrying NMY-2::mKate and NMY-1::GFP knockins (strain FBR241). The acquisition rate was 1 image every 15 minutes, and the playback speed is 6 frames per second.

**Video2**: Two-fold embryo of *nmy-1(mc82[nmy-1::gfp])*control strain. Acquisition rate, 1 frame per second; playback speed, 6 frames per second. Still images of Fig. 3A were taken from this video, with the timing indicated on each image corresponding to the real timing on this videogram.

**Video3**: Double *nmy-2(ne3409ts); nmy-1(mc90ts)::gfp* mutant embryo shifted to 25°C after 2-fold stage. Acquisition rate, 1 frame per second; playback speed, 6 frames per second. Still images of Fig. 3B were taken from this video, with the timing indicated on each image corresponding to the real timing on this videogram.

**Video4**: A *unc-112(RNAi); nmy-1(mc82[nmy-1::gfp])* embryo beyond the 2-fold stage, which had been raised at 25°C. Acquisition rate, 1 frame per second; playback speed, 6 frames per second. Still images of Fig. 3D come from this video, with the timing indicated on each image corresponding to the real timing on this videogram.

**Video5**: A two-fold *nmy-1(mc90ts)::gfp* embryo raised at 25°C. Protein aggregates, embryo rotations and twitching, can be observed. Acquisition rate, 1 frame per second; playback speed, 6 frames, per second.

**Video6**: A 1.5-fold *nmy-2(ne3409ts); nmy-1(mc90ts)::gfp* embryo which had been shifted to 25°C. Note the gradual fusion of the NMY-1::GFP aggregates in the epidermis over time. Acquisition rate, 1 frame per 1 minute; playback speed, 6 frames per second.

## BIBLIOGRAPHY

1. Aigouy, B., R. Farhadifar, D. B. Staple, A. Sagner, J. C. Roper et al., 2010 Cell flow reorients the axis of planar polarity in the wing epithelium of Drosophila. Cell 142: 773–786.

2. Althaus, K., and A. Greinacher, 2009 MYH9-related platelet disorders. Semin Thromb Hemost 35: 189–203.

3. Bonakdar, N., R. Gerum, M. Kuhn, M. Sporrer, A. Lippert et al., 2016 Mechanical plasticity of cells. Nat Mater 15: 1090–1094.

4. Choi, H. J., and W. I. Weis, 2011 Crystal structure of a rigid four-spectrin-repeat fragment of the human desmoplakin plakin domain. J Mol Biol 409: 800–812.

5. Collinet, C., M. Rauzi, P. F. Lenne and T. Lecuit, 2015 Local and tissue-scale forces drive oriented junction growth during tissue extension. Nat Cell Biol 17: 1247–1258.

6. Costa, M., B. W. Draper and J. R. Priess, 1997 The role of actin filaments in patterning the Caenorhabditis elegans cuticle. Dev Biol 184: 373–384.

7. del Rio, A., R. Perez-Jimenez, R. Liu, P. Roca-Cusachs, J. M. Fernandez et al., 2009 Stretching single talin rod molecules activates vinculin binding. Science 323: 638–641.

8. Diogon, M., F. Wissler, S. Quintin, Y. Nagamatsu, S. Sookhareea et al., 2007 The RhoGAP RGA-2 and LET-502/ROCK achieve a balance of actomyosin-dependent forces in C. elegans epidermis to control morphogenesis. Development 134: 2469–2479.

9. Doubrovinski, K., M. Swan, O. Polyakov and E. F. Wieschaus, 2017 Measurement of cortical elasticity in Drosophila melanogaster embryos using ferrofluids. Proc Natl Acad Sci U S A 114: 1051–1056.

10. Gally, C., F. Wissler, H. Zahreddine, S. Quintin, F. Landmann et al., 2009 Myosin II regulation during C. elegans embryonic elongation: LET-502/ROCK, MRCK-1 and PAK-1, three kinases with different roles. Development 136: 3109-3119.

11. Gillard, G., O. Nicolle, T. Brugière, S. Prigent, M. Pinot et al., 2019 Force Transmission between Three Tissues Controls Bipolar Planar Polarity Establishment and Morphogenesis. Curr Biol 29: 1360–1368.e1364.

12. Gilmour, D., M. Rembold and M. Leptin, 2017 From morphogen to morphogenesis and back. Nature 541: 311–320.

13. Goodwin, K., and C. M. Nelson, 2021 Mechanics of Development. Dev Cell 56: 240–250.

14. Guo, S., and K. J. Kemphues, 1996 A non-muscle myosin required for embryonic polarity in Caenorhabditis elegans. Nature 382: 455–458.

15. Hu, X., F. M. Margadant, M. Yao and M. P. Sheetz, 2017 Molecular stretching modulates mechanosensing pathways. Protein Sci 26: 1337–1351.

16. Khalilgharibi, N., J. Fouchard, N. Asadipour, R. Barrientos, M. Duda et al., 2019 Stress relaxation in epithelial monolayers is controlled by the actomyosin cortex. Nature Physics 15: 839-+.

17. Kovacevic, I., J. M. Orozco and E. J. Cram, 2013 Filamin and phospholipase C-ε are required for calcium signaling in the Caenorhabditis elegans spermatheca. PLoS Genet 9: e1003510.

18. Lardennois, A., G. Pasti, T. Ferraro, F. Llense, P. Mahou et al., 2019 An actin-based viscoplastic lock ensures progressive body-axis elongation. Nature 573: 266–270.

19. Law, R., P. Carl, S. Harper, P. Dalhaimer, D. W. Speicher et al., 2003 Cooperativity in forced unfolding of tandem spectrin repeats. Biophys J 84: 533–544.

20. le Duc, Q., Q. Shi, I. Blonk, A. Sonnenberg, N. Wang et al., 2010 Vinculin potentiates E-cadherin mechanosensing and is recruited to actin-anchored sites within adherens junctions in a myosin II-dependent manner. J Cell Biol 189: 1107–1115.

21. Lenne, P. F., A. J. Raae, S. M. Altmann, M. Saraste and J. K. Horber, 2000 States and transitions during forced unfolding of a single spectrin repeat. FEBS Lett 476: 124–128.

22. Levayer, R., A. Pelissier-Monier and T. Lecuit, 2011 Spatial regulation of Dia and Myosin- II by RhoGEF2 controls initiation of E-cadherin endocytosis during epithelial morphogenesis. Nature cell biology 13: 529–540.

23. Liu, J., L. L. Maduzia, M. Shirayama and C. C. Mello, 2010 NMY-2 maintains cellular asymmetry and cell boundaries, and promotes a SRC-dependent asymmetric cell division. Dev Biol 339: 366–373.

24. Lye, C. M., G. B. Blanchard, H. W. Naylor, L. Muresan, J. Huisken et al., 2015 Mechanical Coupling between Endoderm Invagination and Axis Extension in Drosophila. PLoS Biol 13: e1002292.

25. Maitre, J. L., R. Niwayama, H. Turlier, F. Nedelec and T. Hiiragi, 2015 Pulsatile cell- autonomous contractility drives compaction in the mouse embryo. Nat Cell Biol 17: 849–855.

26. Martin, A. C., M. Kaschube and E. F. Wieschaus, 2009 Pulsed contractions of an actin- myosin network drive apical constriction. Nature 457: 495–499.

27. McCullough, B. R., E. E. Grintsevich, C. K. Chen, H. Kang, A. L. Hutchison et al., 2011 Cofilin-linked changes in actin filament flexibility promote severing. Biophys J 101: 151–159.

28. Molnar, K., and M. Labouesse, 2021 The plastic cell: mechanical deformation of cells and tissues. Open Biol 11: 210006.

29. Moore, S. W., P. Roca-Cusachs and M. P. Sheetz, 2010 Stretchy proteins on stretchy substrates: the important elements of integrin-mediated rigidity sensing. Dev Cell 19: 194–206.

30. Munro, E., J. Nance and J. R. Priess, 2004 Cortical flows powered by asymmetrical contraction transport PAR proteins to establish and maintain anterior-posterior polarity in the early C. elegans embryo. Developmental cell 7: 413–424.

31. Niethammer, P., 2021 Components and Mechanisms of Nuclear Mechanotransduction. Annu Rev Cell Dev Biol 37: 233–256.

32. Osorio, D. S., F. Y. Chan, J. Saramago, J. Leite, A. M. Silva et al., 2019 Crosslinking activity of non-muscle myosin II is not sufficient for embryonic cytokinesis in C. elegans. Development 146.

33. Piekny, A. J., J. L. Johnson, G. D. Cham and P. E. Mains, 2003 The *Caenorhabditis elegans* nonmuscle myosin genes *nmy-1* and *nmy-2* function as redundant components of the *let-502*/Rho-binding kinase and *mel-11*/myosin phosphatase pathway during embryonic morphogenesis. Development 130: 5695–5704.

34. Rauzi, M., P. F. Lenne and T. Lecuit, 2010 Planar polarized actomyosin contractile flows control epithelial junction remodelling. Nature 468: 1110–1114.

35. Rogalski, T. M., G. P. Mullen, M. M. Gilbert, B. D. Williams and D. G. Moerman, 2000 The UNC-112 gene in Caenorhabditis elegans encodes a novel component of cell- matrix adhesion structures required for integrin localization in the muscle cell membrane. The Journal of cell biology 150: 253–264.

36. Shelton, C. A., J. C. Carter, G. C. Ellis and B. Bowerman, 1999 The nonmuscle myosin regulatory light chain gene mlc-4 is required for cytokinesis, anterior-posterior polarity, and body morphology during Caenorhabditis elegans embryogenesis. J Cell Biol 146: 439–451.

37. Solon, J., A. Kaya-Copur, J. Colombelli and D. Brunner, 2009 Pulsed forces timed by a ratchet-like mechanism drive directed tissue movement during dorsal closure. Cell 137: 1331–1342.

38. Staddon, M. F., K. E. Cavanaugh, E. M. Munro, M. L. Gardel and S. Banerjee, 2019 Mechanosensitive Junction Remodeling Promotes Robust Epithelial Morphogenesis. Biophys J 117: 1739–1750.

39. Sun, Y., M. Li, D. Zhao, X. Li, C. Yang et al., 2020 Lysosome activity is modulated by multiple longevity pathways and is important for lifespan extension in C. elegans. Elife 9.

40. Vuong-Brender, T. T., M. Ben Amar, J. Pontabry and M. Labouesse, 2017 The interplay of stiffness and force anisotropies drives embryo elongation. Elife 6.

41. Williams, B. D., and R. H. Waterston, 1994 Genes critical for muscle development and function in *Caenorhabditis elegans* identified through lethal mutations. J Cell Biol 124: 475–490.

42. Yang, X., 2017, pp. Université de Strasbourg, France, Strasbourg, France.

43. Yap, A. S., K. Duszyc and V. Viasnoff, 2018 Mechanosensing and Mechanotransduction at Cell-Cell Junctions. Cold Spring Harbor Perspectives in Biology 10.

44. Yonemura, S., Y. Wada, T. Watanabe, A. Nagafuchi and M. Shibata, 2010 alpha-Catenin as a tension transducer that induces adherens junction development. Nat Cell Biol 12: 533–542.

45. Zhang, H., and M. Labouesse, 2010 The making of hemidesmosome structures in vivo. Developmental Dynamics 239: 1465–1476.

46. Zhang, H., F. Landmann, H. Zahreddine, D. Rodriguez, M. Koch et al., 2011 A tension-induced mechanotransduction pathway promotes epithelial morphogenesis. Nature 471: 99–103.

